# PRIMA: a gene-centered, RNA-to-protein method for mapping RNA-protein interactions

**DOI:** 10.1101/074823

**Authors:** Alex M. Tamburino, Ebru Kaymak, Shaleen Shrestha, Amy D. Holdorf, Sean P. Ryder, Albertha J.M. Walhout

## Abstract

Interactions between RNA binding protein (RBP) and mRNAs are critical to post-transcriptional gene regulation. Eukaryotic genomes encode thousands of mRNAs and hundreds of RBPs. However, in contrast to interactions between transcription factors (TFs) and DNA, the interactome between RBPs and RNA has been explored for only a small number of proteins and RNAs. This is largely because the focus has been on using ‘protein-centered’ (RBP-to-RNA) interaction mapping methods that identify the RNAs with which an individual RBP interacts. While powerful, these methods cannot as of yet be applied to the entire RBPome. Moreover, it may be desirable for a researcher to identify the repertoire of RBPs that can interact with an mRNA of interest – in a ‘gene-centered’ manner, yet few such techniques are available. Here, we present Protein-RNA Interaction Mapping Assay (PRIMA) with which an RNA ‘bait’ can be tested versus multiple RBP ‘preys’ in a single experiment. PRIMA is a translation-based assay that examines interactions in the yeast cytoplasm, the cellular location of mRNA translation. We show that PRIMA can be used with small RNA elements, as well as with full-length *Caenorhabditis elegans* 3′UTRs. PRIMA faithfully recapitulates numerous well-characterized RNA-RBP interactions and also identified novel interactions, some of which were confirmed *in vivo*. We envision that PRIMA will provide a complementary tool to expand the depth and scale with which the RNA-RBP interactome can be explored.

## INTRODUCTION

The post-transcriptional regulation of gene expression is vital to organismal development and homeostasis. Post-transcriptional gene regulation affects many aspects of an mRNA, including splicing, 3′-end formation, nuclear-cytoplasmic export, localization, translation and stability (Glisovic et al., 2008). These processes are controlled by physical interactions with different RBPs that often occur through the 3′ untranslated region (UTR)(Moore, 2005; Szostak and Gebauer, 2013).

Thousands of 3′UTRs have been experimentally defined in several model organisms (Derti et al., 2012; Jan et al., 2011; Mangone et al., 2010; Ulitsky et al., 2012). In addition, compendia of hundreds of RBPs encompassing ~5% of all protein-coding genes have been predicted or experimentally determined in various model organisms and humans (Baltz et al., 2012; Castello et al., 2012; Gerstberger et al., 2014; Tamburino et al., 2013). Thus, there is a vast matrix of potential interactions between 3′UTRs and RBPs, or interactomes, that needs to be explored. Several assays are available to identify or study RNA-RBP interactions. Most of these are what we refer to as ‘protein-centered’, or RBP-to-RNA, because they study a single RBP at a time and identify the RNA molecules with which this RBP interacts. These *in vivo* methods include microarray profiling of RNAs associated with immunopurified RBPs (RIP-Chip) (Keene et al., 2006; Tenenbaum et al., 2000), cross-linking of the RBP to the RNA followed by and immunoprecipitation (CLIP) (Ule et al., 2005), plus variations of CLIP that use high-throughput sequencing (HITS-CLIP)(Licatalosi et al., 2008) and Photoactivatable-Ribonucleoside-Enhanced CLIP (Hafner et al., 2010). *In vitro* methods to characterize the binding specificity of RBPs include electromobility shift (EMSA) and RNAcompete assays, which can be used to test binding of individual RBPs to single or multiple RNA elements, respectively (Pagano et al., 2011; Ray et al., 2009). These methods can be limited in their use because they require suitable anti-RBP antibodies or purified RBPs, because they are carried out *in vitro*, or because they cannot be used in a gene-centered, or RNA-to-RBP manner, which is what one would like to do when the focus is a single gene, an individual 3′UTR, or a particular RNA element or structure.

Several RNA-to-RBP interaction mapping methods have been developed, including proteomic methods that involve the pull-down of mRNAs or non-coding RNAs using oligo d(T) beads (Butter et al., 2009; Castello et al., 2012; Matia-Gonzalez et al., 2015), and examining the precipitated RBP interactome by mass spectrometry. This type of approach identifies tens to hundreds of putative RBPs, but provides no information about whether the interaction is direct or indirect, or if it is specific to a particular structure or sequence. Further, these approaches can be challenging to apply to intact organisms or tissues due to cellular heterogeneity and (low) RBP or mRNA expression levels. A heterologous method that can be used in either an RBP-to-RNA or RNA-to-RBP configuration is the yeast three-hybrid (Y3H) system. This system is based on the reconstitution of a functional transcription factor via an RNA-RBP interaction in nucleus of yeast cells (SenGupta et al., 1996). However, many RNA-RBP interactions occur in the cytoplasm. Further, Y3H assays can be limited by the length and nucleotide sequence of the RNA (Zhang et al., 1999).

The nematode *Caenorhabditis elegans* is a powerful model organism for the study of biological interactome networks (Lee et al., 2008; Li et al., 2004; MacNeil et al., 2015; Reece-Hoyes et al., 2013; Walhout et al., 2000a). *C. elegans* transgenic strains can be generated that express a fluorescent reporter protein under the control of a promoter (with fixed 3′UTR)(Chalfie et al., 1994; Grove et al., 2009; Hunt-Newbury et al., 2007; Martinez et al., 2008; Ritter et al., 2013), or 3'UTR (with fixed promoter) of interest (Merritt et al., 2008). Such strains can then be used with RNAi knockdown screening to identify or characterize proteins that regulate that promoter or 3′UTR either directly or indirectly (MacNeil et al., 2015; Watson et al., 2013)((Merritt et al., 2008)). We predicted that the *C. elegans* genome contains up to 887 RBPs, and this estimate has largely been verified by proteomic findings (Matia-Gonzalez et al., 2015; Tamburino et al., 2013). *In vitro* assays have been used to determine the binding specificities of several *C. elegans* RBPs (Farley et al., 2008; Pagano et al., 2011; Pagano et al., 2007). However, it has proven difficult to use these specificities to predict complex mRNAs that are bound by the RBP and, therefore, RBP interactions with larger mRNA 3′UTRs remain largely unexplored. Most studies of RBPs in *C. elegans* have been limited to protein-centered methods, examining RNA targets of specific RBPs, including Y3H studies (Bernstein et al., 2005; Koh et al., 2009; Opperman et al., 2005; Stumpf et al., 2008). To our knowledge, RNA-centered studies have been limited to a few *in vitro* yeast-based assays and one proteomics study (Hook et al., 2005; Matia-Gonzalez et al., 2015), illustrating the need for additional methods and tools.

We have shown extensively that the mapping of the transcription factor interactome greatly benefits from the use of multiple complementary approaches, including both protein- and DNA-centered methods (Fuxman Bass et al., 2015; Reece-Hoyes et al., 2013). Multiple, complementary methods are needed to map networks because not all proteins are amenable to protein-centered methods, because experiments with intact organisms have different caveats, and because no single method will be able to capture the entire interactome (Walhout, 2011).

Here, we present PRIMA, a gene-centered <u>P</u>rotein-<u>R</u>NA <u>I</u>nteraction <u>M</u>apping <u>A</u>ssay that can be used to study RNA-RBP interactions with a variety of RNA elements or 3′UTRs, and different RBPs within the cytoplasm of yeast cells, the cellular milieu where many RBP-RNA interactions occur. PRIMA enables the pairwise testing of numerous RBPs for their capacity to bind an RNA of interest in a single experiment. PRIMA is based on the stabilizing effect of a physical interaction between the 3′ end and 5′ end of an mRNA, which results in effective translation. PRIMA uses expression of the green fluorescent protein (GFP) as a reporter. The fluorescent signal is detected in a quantitative manner using high-throughput flow cytometry, and positive interactions are calculated using computational data processing and statistical analyses of replicates. We show that PRIMA can be used with small RNA elements, as well as with *C. elegans* 3′UTRs to capture known and novel interacting RBPs. PRIMA will provide an addition to the toolkit for the mapping of the RNA-RBP interactome.

## DESIGN

PRIMA is based on the endogenous function of yeast poly(A)-binding protein (Pab1p), which binds the 3′ poly(A) tail and interacts with the 5′ end of an mRNA through the scaffold protein, eIF4G, and the cap binding protein, eIF4E, thereby stabilizing the mRNA and increasing translation of the mRNA into protein (Mangus et al., 2003). We reasoned that we could reconstitute this interaction by using a reporter mRNA that encodes GFP and replacing its poly(A) tail with a selected RNA ‘bait’ element (*e.g.*, a 3′UTR) of interest, and fusing a candidate interacting ‘prey’ RBP to Pab1p (Figure 1A). When the RBP binds the RNA element, Pab1p interacts with the 5′ end of the reporter mRNA resulting in stabilization and production of GFP. However, when challenged with a non-interacting RBP, the mRNA is unstable and little GFP is produced.

**Figure 1.**
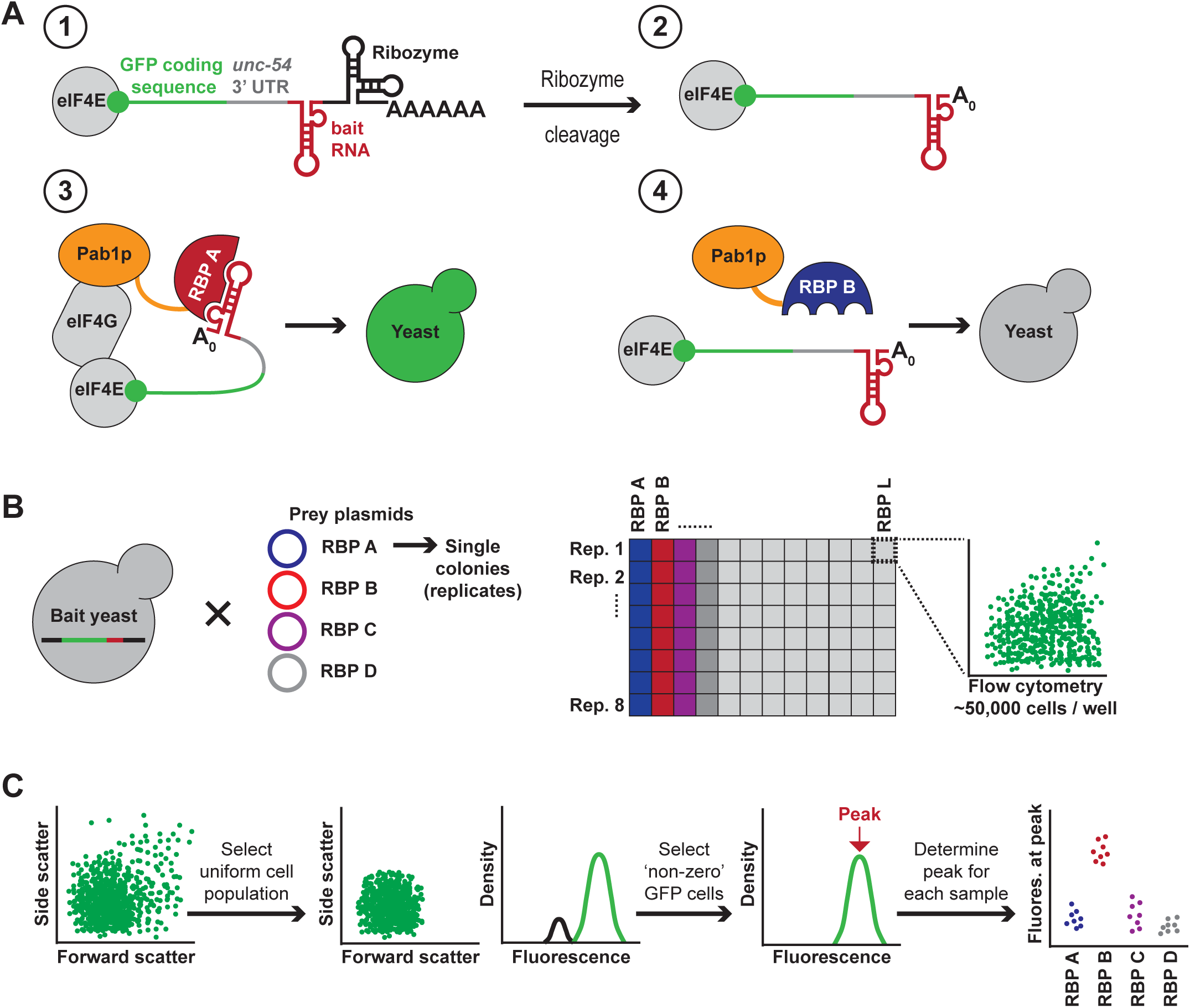
PRIMA Design and Experimental Workflow. (A) In PRIMA, RNA-RBP interactions are measured by GFP expression from a reporter mRNA or ‘RNA bait’. RBP ‘preys’ are fused to Pab1p, which binds the translation initiation machinery when bound to the 3′ end of the mRNA. The GFP reporter mRNA (green) including a minimal *unc-54* 3′UTR (gray) and an RNA bait (red) is expressed without a poly(A) tail by using a *cis*-encoded, self-cleaving hammerhead ribozyme (black) (part 1). An RBP-Pab1p fusion protein (red or blue) is co-expressed with the reporter bait RNA. When the RBP binds the RNA element of interest, the mRNA is stabilized and translated resulting in increased GFP levels (part 3). In contrast, when the bait mRNA and RBP prey do not interact the mRNA is unstable and the GFP signals remain low (part 4). (B) A yeast RNA bait strain is transformed with an RBP-Pab1p-encoding plasmid. Multiple plasmids can be transformed in parallel. Independent colonies are isolated and grown to log phase in liquid media. GFP expression is measured in ~50,000 cells per replicate using automated flow cytometry. (C) Data filtering. The 50% most uniform cells are selected according the forward scatter (FSC, size) and side scatter (SSC, granularity) dot plot profiles. Next, fluorescence of the uniform cells is plotted as a Kernel density plot and ‘non-zero’ GFP positive cells are selected to ensure basal mRNA expression. The minimum fluorescence threshold (FL1>2048 *i.e.* fluorescence) is determined using GFP(-) control cell populations. Finally, the peak fluorescence was determined for each replicate (see Experimental Procedures for details).

To avoid endogenous Pab1p from binding to and stabilizing the reporter mRNA, we removed the poly(A) tail by adding a *cis*-encoded, self-cleaving hammerhead ribozyme (Dower et al., 2004) to the 3′ end of the mRNA, just 5′ of the poly(A) tail (Figure 1A). Ribozyme cleavage removes the 3′ end of the message, leaving it unable to be protected from degradation by Pab1p. Finally, we added a generic *C. elegans unc-54* 3′UTR upstream of the RNA bait/ribozyme and downstream of the GFP-encoding open reading frame to facilitate RNA export to the cytoplasm (Dower et al., 2004; Okkema et al., 1993).

The first step in a PRIMA experiment is to generate a yeast bait strain that produces the reporter mRNA in which the RNA element of interest is located in between the *unc-54* 3′UTR and the ribozyme (Figure 1A). The second step involves the transformation of the RNA bait strain with a plasmid encoding a chimeric protein consisting of an RBP and Pab1p. GFP expression is then measured in ~50,000 cells per transformant, using automated flow cytometry (Figure 1B). Once collected, the data is filtered to select cells of uniform size and morphology. Next, ‘non-zero’ fluorescent cells are selected and the peak density of the population is calculated for each replicate (Figure 1C and **S1A**, **S1B**). The peak density is then compared across the dataset to determine positive RNA-RBP interactions.

## RESULTS

### Detection of Known RNA-RBP Interactions

As a proof-of-concept we used two well-characterized RNA-RBP interactions: one involving the bacteriophage MS2 stem-loop binding site (MS2BS), which interacts with the MS2 coat protein (MS2), and the other being the stem-loop binding element from the 3′ end of histone mRNAs (HBE) that binds the mammalian stem-loop binding protein (SLBP)(Johansson et al., 1998; Michel et al., 2000). We tested each RNA bait versus both RBPs to simultaneously assess PRIMA's sensitivity and specificity. Quantification by flow cytometry showed that PRIMA could detect each test interaction with high specificity as only the cognate pairs activated GFP expression (Figure 2A and 2B). Perhaps not surprisingly, there is a spread of fluorescence between the individual strains, indicating the need for multiple replicates and statistical testing.

**Figure 2.**
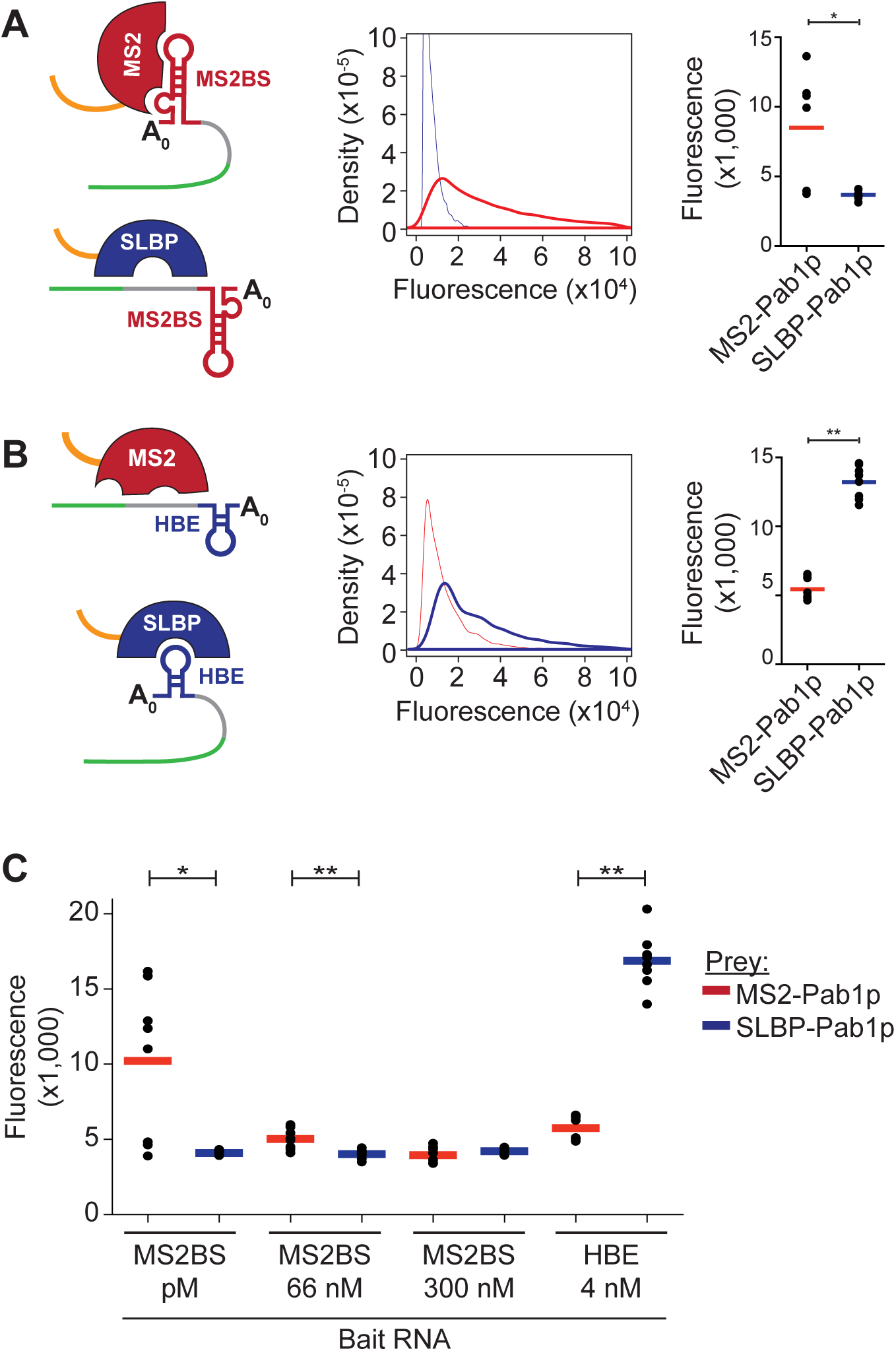
PRIMA Validation. (A) The MS2BS stem-loop RNA bait was tested with its known RBP partner MS2 and a non-binding RBP SLBP. Kernel density plot vs. GFP fluorescence: positive interaction (red curve) and negative control interaction (blue curve). Dot plots show the peak fluorescence for each of the eight replicates. The bar represents the mean of eight independent replicates. (**p<0.01, *p<0.05, student's t-test). (B) The same experiment as Part A, only the HBE stem-loop is the RNA bait with its partner SLBP, while the MS2 RBP is the negative control. Kernel density plot vs. GFP fluorescence: positive interaction (blue curve) and negative control interaction (red curve). (C) High (MS2BS pM) and medium (MS2BS 66 nM) RNA-RPB affinity interactions can be detected by PRIMA for the MS2 RBP, while low affinity (MS2BS 300 nM) and non-specific (HBE 4nM) interactions cannot be detected. The bar represents the mean of eight independent replicates. (**p<0.01, *p<0.05, student's t-test).

We further assessed the sensitivity of PRIMA by introducing two different single nucleotide point mutations in the MS2BS that reduce the interaction affinity of MS2 to 66 nM and 300 nM, respectively (Johansson et al., 1998). As expected, the highest degree of GFP expression occurs with the original, high-affinity MS2BS (pM affinity). The 66 nM interaction moderately induced GFP expression yet still showed a statistically significant difference between prey interactions, while the low-affinity interaction (300 nM) was not detected by PRIMA (Figure 2C). In all cases the MS2BS showed no significant fluorescence with the SLBP-Pab1p prey. Thus, PRIMA can detect specific interactions with native RNAs and their cognate RBPs.

### Optimizing PRIMA

We tested several known interactions with *C. elegans* RBPs (Figure 3A). Initial attempts failed to specifically induce high levels of GFP expression in any of the test cases (**Figure S2A**). There are several potential reasons for low sensitivity, including poor expression of the bait mRNA reporter or RBP prey in yeast, mislocalization of the prey, for instance to the nucleus, or toxic effects of prey expression. To address these issues, we first introduced a high-affinity MS2BS to the 3′ end of each RNA bait (Figure 3B). This modification allowed us to determine that the RNA baits used are functional in PRIMA because co-expression with MS2-Pab1p increased GFP expression for all baits tested (**Figure S2B**). Second, we tested whether any of the RBP preys were toxic to yeast. We obtained no or very few colonies upon transformation of the GLD-1-encoding plasmid, suggesting that expression of this RBP is toxic to yeast (**Figure S2C**). Third, we tested the functionality of the other preys by expressing them as RBP-MS2-Pab1p fusion proteins and introducing these constructs into the bait strain harboring a GFP reporter with a high-affinity MS2BS as RNA bait (**Figure S2D**). GFP was induced by all five of the *C. elegans* RBP-MS2-Pab1p preys tested, demonstrating that all RBPs are appropriately expressed and localized. Altogether, these results indicate that, with the exception of the one toxic RBP, all baits and preys tested are functional within the context of PRIMA. Therefore, we hypothesized that the cognate RBP-mRNA interaction affinities may be below the detection limits of PRIMA.

**Figure 3.**
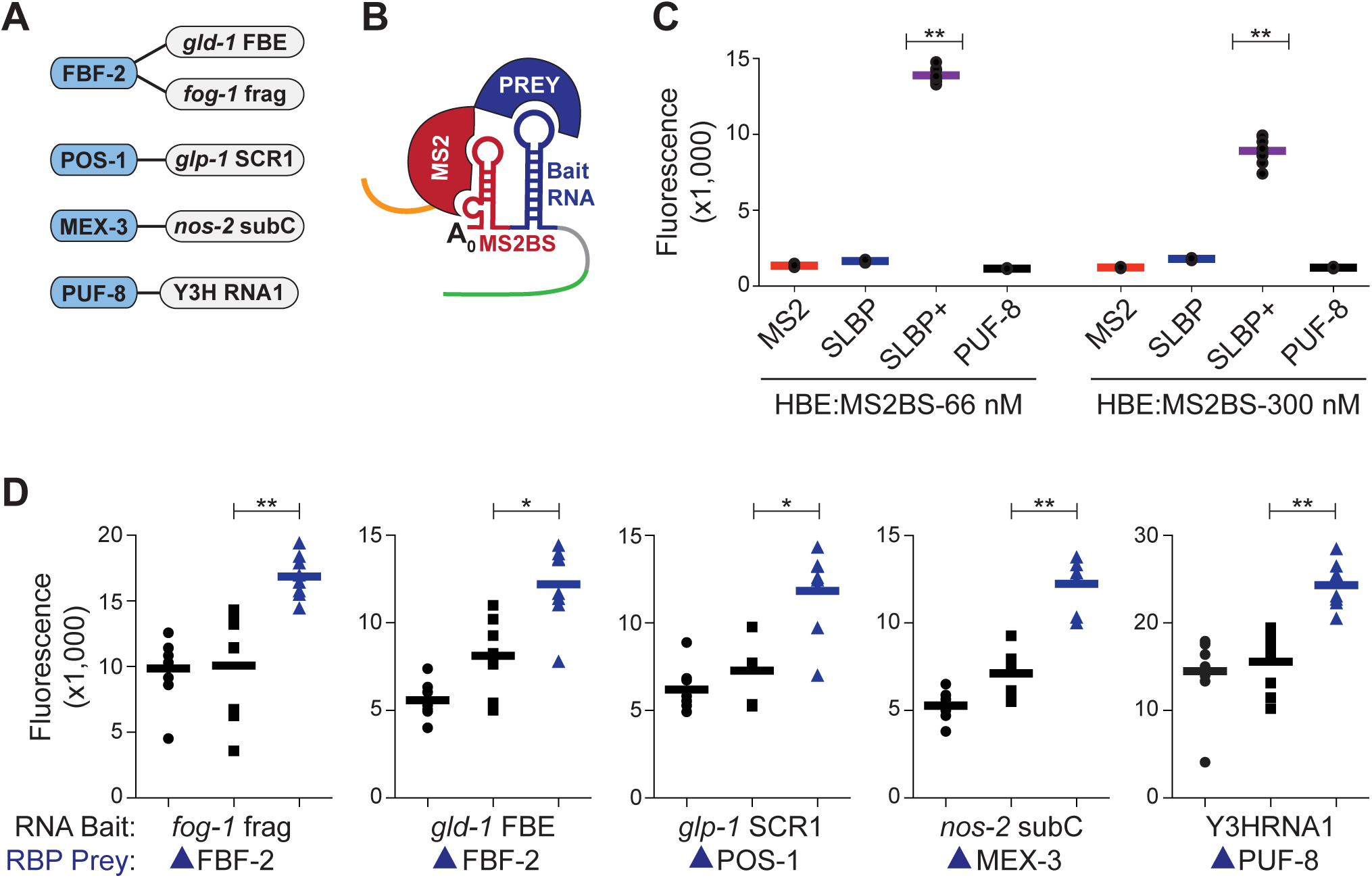
Known RNA-RBP interactions can be detected by PRIMA. (A) RNA Binding Domains (blue) were tested for interactions with their known RNA elements (white). (B) Schematic of the modified bait strain (green, GFP; grey, 3′UTR; blue, bait RNA; red, weak affinity MS2BS; blue half circle, Prey RBP; red half circle, MS2 RBP; orange, Pab1p). (C) Fusion baits containing both HBE and weak and low affinity MS2BS were tested against single RBP-Pab1p preys and SLBP-MS2-Pab1p (SLBP+) prey as a proof-of-concept. PUF-8-Pab1p is included as a non-binding negative control. (**p<0.001, student's t-test). (D) Fluorescence levels for each RNA-RBD interaction. SLBP-Pab1p (•) and SLBP-MS2-Pab1p (■) preys were negative controls for each bait. Bars indicate the mean fluorescence for all eight replicates. Positive interactions are shown in blue (*p<0.01, **p<0.001, student's t-test).

We reasoned that the sensitivity of PRIMA could be improved by including a high specificity, low-affinity driver interaction adjacent to the test interaction. We selected the interaction between MS2BS and MS2 because it is highly specific, and it can be modified to lower affinities. We introduced the moderate (66 nM) or low-affinity (300 nM) MS2BS at the 3′ end of each RNA bait (Figure 3B). Additionally, we added the MS2 protein to the preys to create RBP-MS2-Pab1p fusion proteins. To test whether these modifications result in enhanced sensitivity, we used the SLBP prey, and found that GFP production was dramatically increased when the SLBP-MS2-Pab1p prey was tested with RNA baits that are located adjacent to either a moderate or low-affinity MS2BS (Figure 3C).

Next, we re-assayed the test set of RNA-RBP interactions using the MS2 fusion strategy. The 300 nM low affinity MS2BS was fused to each RNA bait because this sequence show little background binding in the presence of MS2-fused RBPs (Figure 3D). RNA-binding domains (RBD) were used in place of full-length RBPs to reduce potentials for steric hindrance. Additionally, bait constructs were integrated into the yeast genome to reduce cell-to-cell variability in bait RNA expression. Five RNA baits were tested against four RBD preys (Figure 3A). These preys contain different types of RBDs: FBF-2 and PUF-8 contain PUF domains, MEX-3 has a KH domain, and POS-1 contains CCCH zing finger. SLBP-Pab1p and SLBP-MS2-Pab1p were included as negative controls for basal GFP expression and increases mediated by MS2 binding, respectively. Previously characterized interactions were detected for all five RNA baits (Figure 3D). Two of these, *fog-1* fragment and *gld-1* FBF binding element (FBE), were bound by FBF-2 as expected (Bernstein et al., 2005; Thompson et al., 2005). The *glp-1* SCR1 was bound by POS-1 (Farley et al., 2008; Farley and Ryder, 2012). The *nos-2* subC fragment was bound by MEX-3 (Jadhav et al., 2008; Pagano et al., 2009). The previously characterized Y3HRNA1 fragment interaction with PUF-8 was also confirmed by PRIMA (Opperman et al., 2005). Overall this reference set demonstrates that PRIMA can detect previously known *C. elegans* RNA-RBP interactions involving different types of RBDs.

### PRIMA Can Use Full Length 3′UTRs as Bait

Next, we asked whether PRIMA can detect RNA–RBP interactions with full-length 3′UTRs as RNA baits. We selected six *C. elegans* 3′UTRs: *nos-2* (318 nt) and *glp-1* (363 nt), *mex-3* (437 nt), *atg-4.2* (104 nt), *set-6* (284 nt) and *usp-14* (213 nt), and tested these versus a mini-library of 40 *C. elegans* prey RBPs that are expressed in the germline (Tamburino et al., 2013; Wang et al., 2009). These included several well-characterized RBPs such as POS-1 (which binds *glp-1* and *mex-3*)(Farley et al., 2008; Farley and Ryder, 2012; Ogura et al., 2003), MEX-3, which binds and regulates *glp-1* and *nos-2* (Pagano et al., 2009) and PUF-5, which binds and regulates *glp-1* (Hubstenberger et al., 2012; Lublin and Evans, 2007). For each 3′UTR, PRIMA detected several RBP preys that significantly activated GFP expression (Figure 4).

**Figure 4.**
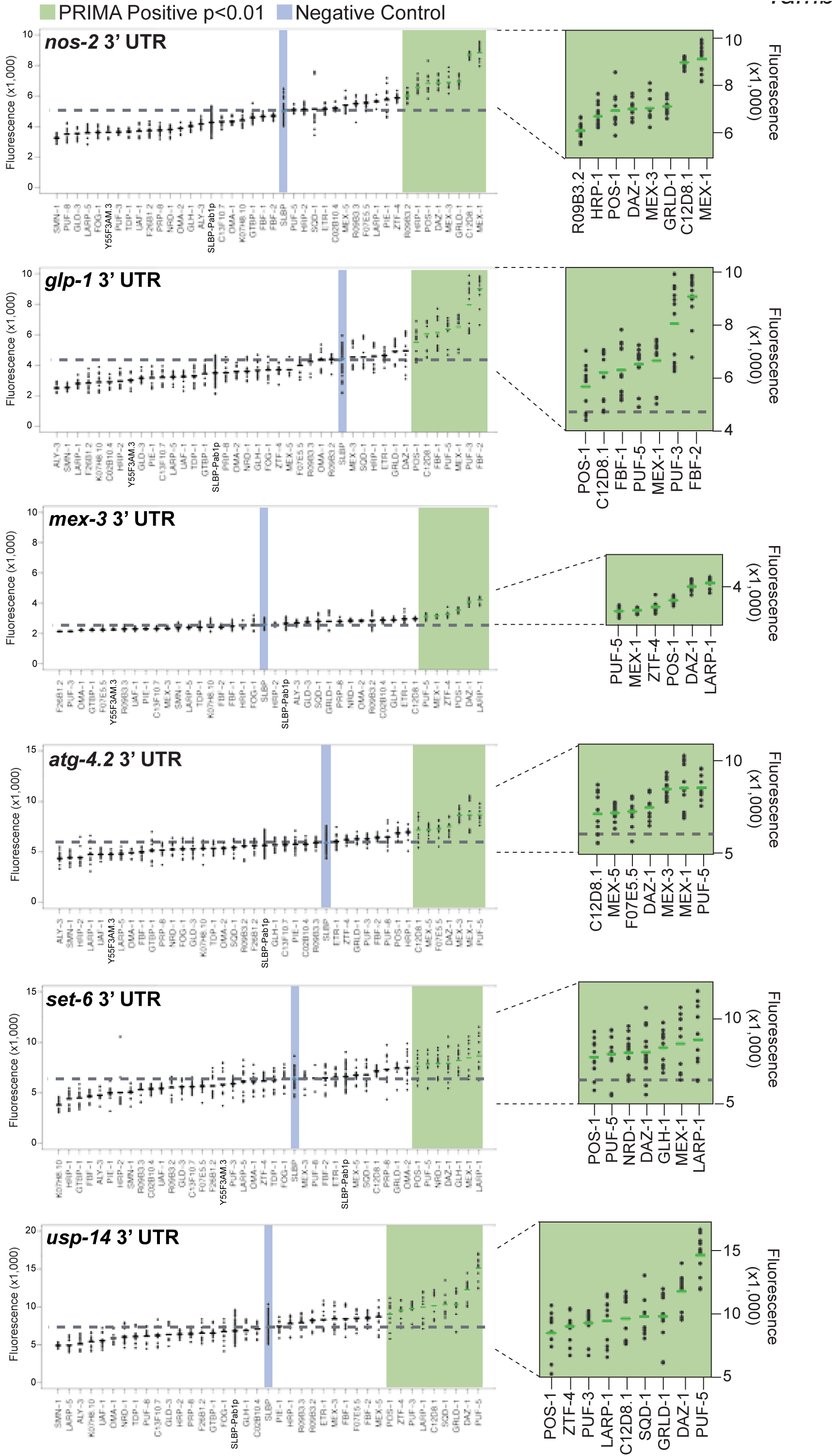
Identification of known and novel *C. elegans* RNA-RBP interactions using full-length 3′UTRs and a RBP prey mini-library. Specific interacting RBPs were detected for six full-length 3′UTRs. Two sets of eight biological replicates were measured for each prey. The fluorescence intensity at the peak was measured for each and the two highest and two lowest samples were removed. The remaining 12 replicates were plotted and the average intensity for each prey is shown. Preys with average intensity >1.20 fold compared to negative control are shown in green (p<0.01, student's t-test). Preys are labeled on the x-axis and include the fusion of MS2 to the prey (except for SLBP-Pab1p).

These interactions are visualized in network format in Figure 5. From these data, we can glean, for the first time, differences between 3′UTRs as well as RBPs using data obtained from a single experiment that was carried out under exactly the same conditions. First, interacting RBPs were identified for each 3′UTR and the number of interacting proteins ranged from six for *mex-3*, to nine for *usp-14*. Secondly, we detected interactions for half of the 40 RBPs tested, most of which have not been studied for their RNA binding specificity prior to our study. Half of the detected RBPs bind only one of the 3′UTRs tested, while four bound five of the six 3′UTRs. These data indicate that PRIMA can detect specific interactions, both for 3′UTRs, and for RBPs, and, with increasingly comprehensive RBP libraries, has the potential to greatly expand the knowledge of the RNA-RBP interactome.

**Figure 5.**
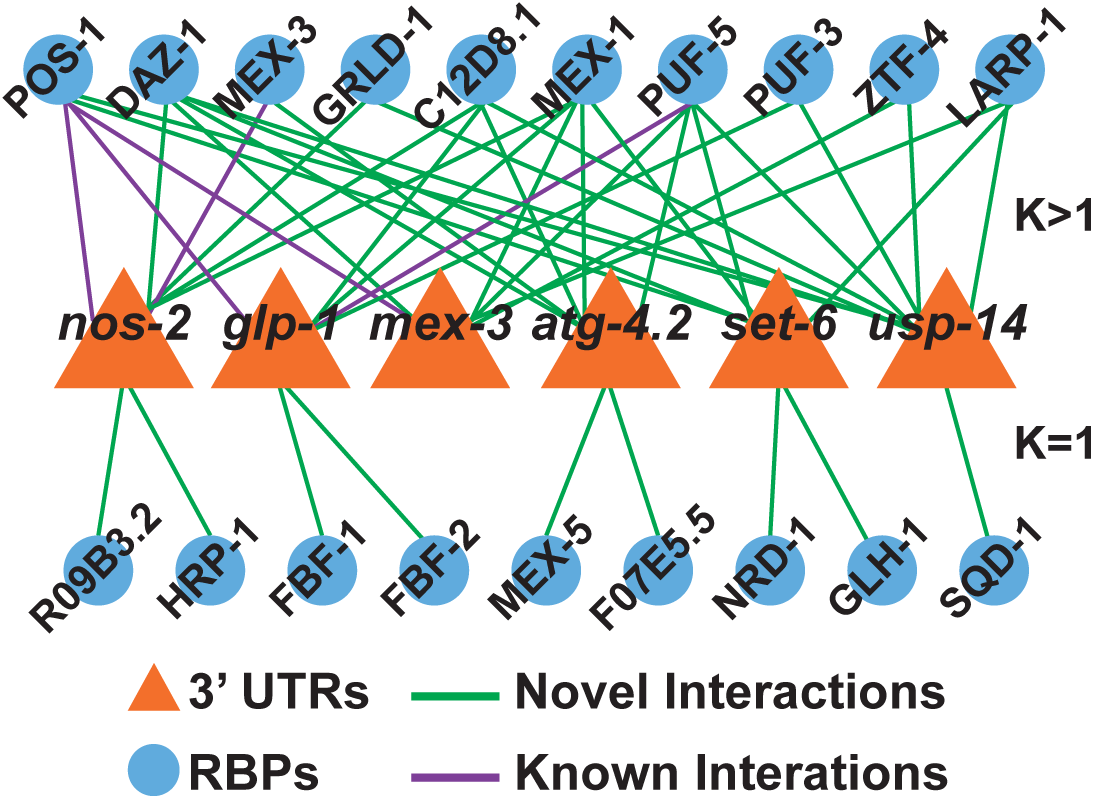
Network graph of known and novel RNA-RBP interactions detected by PRIMA

### PRIMA Can Detect Biologically Active Interactions

The 3′UTRs and RBPs tested are all expressed in the *C. elegans* germline (cartoon in Figure 6A). We used RNAi knockdown of five RBPs that interact with the *glp-1* 3′UTR in PRIMA (Figure 6B), using single copy transgenic animals that express labile GFP under the control of the *mex-5* promoter, which is broadly active in the *C. elegans* germline (Merritt et al., 2008), and under the control of the *glp-1* 3′UTR, which restricts expression to the distal end of the germline (Farley and Ryder, 2012; Pagano et al., 2009). As previously reported, GFP levels increased in the posterior cells of the 4-cell stage embryo of the *glp-1* 3′UTR strain following RNAi-mediated knockdown of *pos-1* (Farley and Ryder, 2012) (**Figure S3**). Importantly, GFP levels also increased in the developing oocytes following RNAi of either *puf-3* or *puf-5* (Figure 6B, 6C). While *puf-5* was known to regulate *glp-1* (Lublin and Evans, 2007), the interaction with *puf-3* is novel. Altogether, these results indicate that PRIMA can detect biologically relevant interactions.

**Figure 6.**
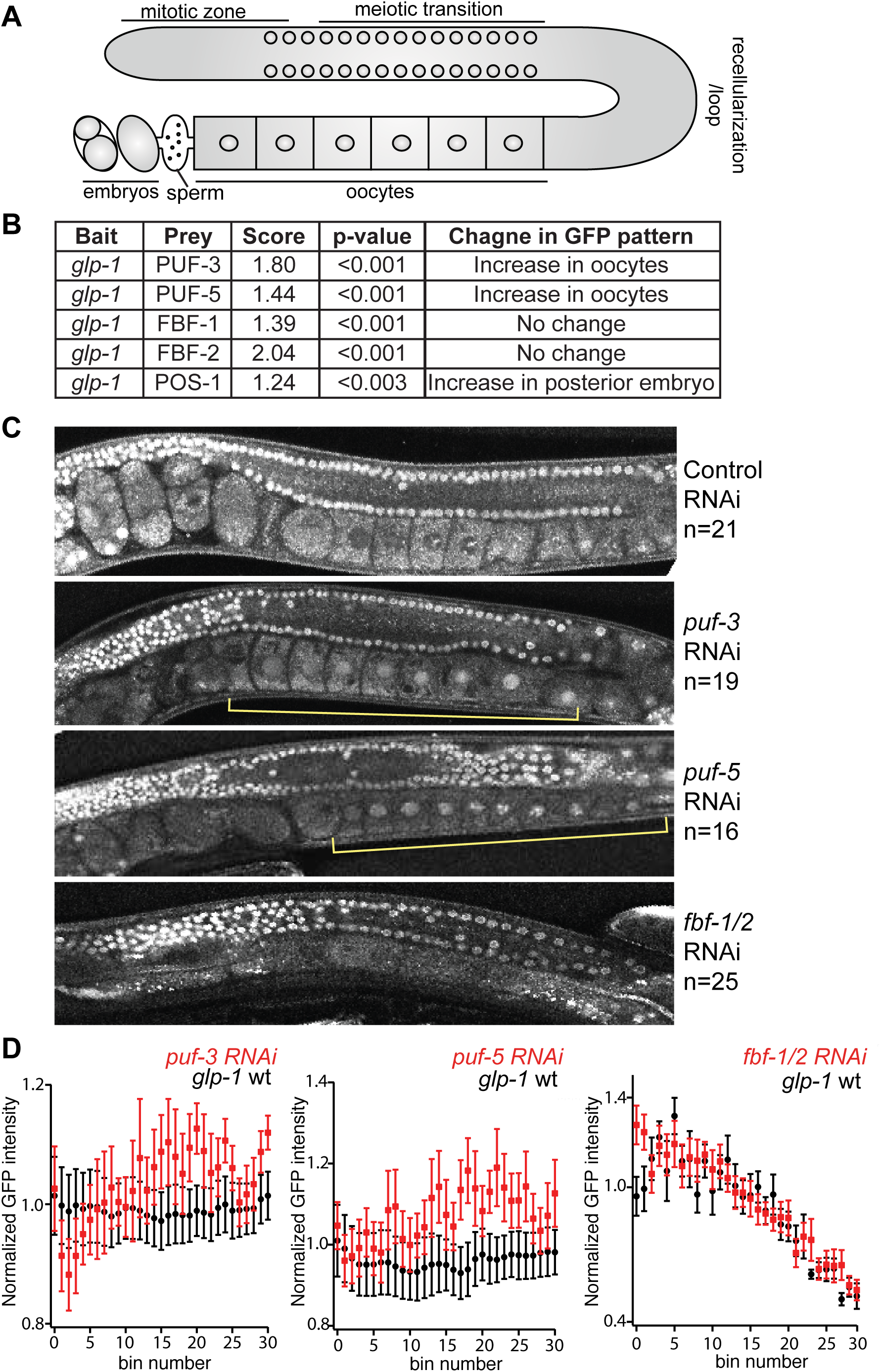
*In vivo* validation of interactions involving RBPs that bind the *glp-1* 3'UTR. (A) Schematic of the *C. elegans* germline. The syncytial region of nuclei is shown in the distal arm of the gonad. The oocytes and the embryos are shown in the proximal are of the gonad. (B) Five RBPs found to interact with the *glp-1* 3′UTR were tested by RNAi *in vivo*. (C) The GFP expression patterns of single copy integrated GFP reporter strains that express GFP under the control of the *glp-1* 3′UTR is shown in the top image. The expression level throughout the germline of the reporter fusion treated with control RNAi is compared to the expression pattern of the strain treated with RNAi to *puf-3*, *puf-5*, and *fbf-1;fbf-2*. Yellow bars denote a change in expression levels in oocytes observed under *puf-3* and *puf-5* RNAi conditions. (D) Quantifications of the confocal images of the *glp-1* reporter strains under the RNAi conditions described above. GFP intensities normalized to average pixel intensity of wild-type oocytes are plotted against bin-number. Red plots show intensities measured under RNAi treatment conditions whereas black bars show intensities measured under control conditions.

## DISCUSSION

PRIMA provides a novel protein-RNA interaction mapping assay that can be used to identify and study RBPs that interact with an RNA element or 3′UTR of interest. We have focused the testing of PRIMA using *C. elegans* RNAs and RBPs, although the method should be applicable to interactions from a variety of organisms.

To our knowledge very few RNA-RBP interactions have been examined in *C. elegans*, and most of these prior studies have been protein-centered to identify RNAs associated with an RBP of interest, or yeast three-hybrid analysis (Table 1). One group has studied RNA-RBP interactions on a proteomic level in *C. elegans* mixed stage and L4 animals, using oligo(dT)_25_ beads followed by mass spectrometry analysis, and identified 549 RBPs (Matia-Gonzalez et al., 2015). However, it is not clear whether these RBPs bind to specific RNA sequences or structures, if some of the co-precipitate with other RBPs.

**Table 1.** Comparison of RNA-RBP interaction detection methods. Assay directionality, advantages and disadvantages of each method, and how often they are used to study *C. elegans* RBPs.

| Method | Directionality | Advantages | Disadvantages | Used to study *C. elegans* RBPs? | Number of *C. elegans* publications |
| --- | --- | --- | --- | --- | --- |
| PRIMA | RNA-Centered | Test specific RNAs, full-length RNA baits, cytoplasmic milieu, no need for antibodies or protein purification | *in yeast*, requires RBP library, cannot detect dimers or modified proteins | Yes | This study |
| Yeast three-hybrid | RNA- or Protein-Centered | Bi-directional, no need for antibodies or protein purification | *in yeast*, nuclear milieu, short RNA baits (<150 nt), requires RBP library, cannot detect dimers or modified proteins | Yes | Protein-centered: ~20, RNA-centered: <5 |
| RNA Interaction Capture | RNA-Centered | *in vivo*, capture entire RBP-ome bound by all polyadenylated RNAs | No specificity, need very large amount of starting material, need RNA-capture beads for pull down, challenging to detect low abundance interactions | Yes | 1 |
| Electrophoretic Mobility Shift Assay (EMSA) | Protein-Centered | Quantitative RBP binding site determination | Requires purified RBPs, low throughput, binding specificity has limited predictive power | Yes | >40 |
| RAP-MS (RNA Antisense Purification) | lncRNA-centered | *in vivo*, genome-scale, quantitative | Limited UV crosslinking efficiency, need large amount of starting material, challenging to detect low abundance interactions | No | 0 |
| ChiRP-MS (Chromatin Isolation by RNA Precipitation) | lncRNA-centered | *in vivo*, genome-scale, quantitative | No specificity, may be difficult to design probe, need a large amount of starting material | No | 0 |
| RIP/RIP-Chip (RBP immunoprecipitation-microarray) | Protein-centered | Physiological (native) conditions, genome-scale RNA detection, *in vivo* | Requires high affinity antibodies or epitope tagged RBPs, post-lysis *in vitro* association of RBPs with spurious targets, lots of contaminating rRNA | Yes | 3 |
| CLIP (including PAR-CLIP and HITS-CLIP) | Protein-centered | Cleaner than native conditions, genome-scale RNA detection, *in vivo*, can determine precise RBP binding site on RNA | Non-physiologic conditions, Requires high affinity antibodies or epitope tagged RBPs, low efficiency of cross-linking | Yes | 1 |
| RNAcompete | Protein-centered | High-throughput binding site determination | Requires purified RBPs, short RNA libraries, binding specificity has limited predictive power | Yes | Some *C. elegans* proteins were included |

PRIMA will provide a gene-centered method to the expanding toolkit for mapping RBP-RNA interactions. It is important to note that PRIMA, like any method, has different advantages and disadvantages (Table 1), and therefore should be thought of as complementary to other techniques. Advantages of PRIMA, aside from being gene-centered, include its ability to use relatively long RNA fragments as bait. For instance, while the Y3H system is limited to 150 nucleotide baits (Zhang et al., 1999) we have shown that fragments nearly three times the length (the *mex-3* 3′UTR, which is 437 nucleotides long) can be used effectively. An additional advantage of PRIMA is that it does not require anti-RBP antibodies, the purification of large numbers of proteins, or a large number of animals to detect interactions. This advantage will likely enable studying RBPs that were heretofore not amenable to interactome studies.

Finally, it is important to note that not all RNA-RBP interactions detected by PRIMA may be biologically meaningful. Indeed, more evidence is becoming available that not all physical transcription factor-DNA interactions, detected either *in vivo* or by yeast-based methods, have a (measurable) regulatory consequence *in vivo* (Kemmeren et al., 2014; MacNeil et al., 2015; Walhout, 2011). This finding could be because the potential regulatory effects were examined under irrelevant physiological conditions, because the interaction effect is masked by redundantly functioning RBPs, or because the interaction is harmless, and can occur without any regulatory consequence (and thus would not be selected for or against).

### Limitations

PRIMA may not detect low affinity RNA-RBP interactions and therefore may miss some important RBPs (Table 1). The addition of the MS2 coat protein at the 5′ end of the RBP prey may sterically hinder some RBP prey-RNA bait interactions. As PRIMA is a yeast-based assay, it does not detect *in vivo* interactions that may lead to problems such as poor expression in yeast or competition with endogenous yeast proteins. Further, RNA-RBP interactions that depend on post-translational modifications of the RBP, or on protein co-factors, will not be detected. However, our successful use of yeast one-hybrid (Y1H) assays for assessing transcription factor (TF)-DNA interactions demonstrates that this type of approach is extremely useful despite such limitations (Deplancke et al., 2006; Fuxman Bass et al., 2015; Reece-Hoyes et al., 2013). The *C. elegans* RBP library is currently small with 40 RBPs, but we anticipate expanding this library as we have done previously for our transcription factor collection (Reece-Hoyes et al., 2011). In the future, we also anticipate streamlining the PRIMA pipeline such that we can make the process higher throughput, similar to yeast one-and-two hybrid assays used for the study of protein-DNA and protein-protein interactions, respectively (Reece-Hoyes et al., 2011; Yu et al., 2011). We have not tested 3′UTRs longer than 437 nucleotides. It is important to note that a majority of 3′UTRs in *C. elegans* are shorter (Mangone et al., 2010), indicating that PRIMA should be broadly applicable to this organism's RNA-RBP interactome. However, human 3′UTRs are on average longer and are frequently alternatively polyadenylated (Derti et al., 2012). We envision that the future development of PRIMA-compatible RBP libraries from different organisms, together with the cloning of full-length 3′UTRs will enable the broad and deep exploration of the RNA-protein interactome, which is essential to gain systems-level insights into post-transcriptional gene regulation.

## EXPERIMENTAL PROCEDURES

### Cloning of RNA Elements and RBPs

All DNA sequences and plasmid configurations used in this manuscript are available in **Table S1** and **Figure S4**. The 3′UTR sequences were taken from the worm UTRome (http://tomato.biodesign.asu.edu/cgi-bin/UTRome/utrome.cgi)(Blazie et al., 2015).

The pADH1::GFP:*unc-54*:MCS:Ribozyme plasmid expression vector was generated using sequential PCR stitching and gap repair of DNA constructs(Orr-Weaver et al., 1983) into the pDest22 backbone (Life Technologies). The S65T GFP sequence was amplified from pFA6:GFP (kindly provided by Paul Kaufman). The shortest *unc-54* 3′UTR isoform is included in all RNA baits. It was amplified from the 3′UTRome entry vector (Mangone et al., 2010). The multiple cloning site (MCS) and hammerhead ribozyme were generated synthetically (Life Technologies). Binding sites were inserted into the MCS of the expression vector using yeast gap repair of synthetic oligos into *Afl*II/*Sma*I (NEB) or *Afl*II/*Cla*I (NEB) digested vectors.

The pGPD:eGFP:*unc-54*:HBE:Stem-loop:Ribozyme integration expression vector was generated from pAG303GPD-EGFP-ccdB (Alberti et al., 2007) by inserting the 3′ end of pADH1:GFP:*unc-54*:HBE:Stem-loop:Ribozyme vector (this work) into the *Not*I/*Sal*I (NEB) fragment. Additional RNA element constructs were generated by replacing the *Afl*II/*Cla*I fragment with synthetic oligos. 3′UTR constructs were generated by replacing the *Eco*RI/*Cl*aI fragment with PCR products amplified from *C. elegans* cDNA.

The pDest Pab1p vector was generated using a Gateway cassette PCR product amplified from pGBKCg (Stellberger et al., 2010) using Platinum HiFi Taq (Invitrogen) and TA cloned into pGEMT (Promega). The *Sac*II/*Xho*I digested product was ligated into the *Sac*II/*Xho*I site of YCplac111-MS2–Pab1p (Amrani et al., 2004) (kindly provided by Allan Jacobson). The pDest-MS2-Pab1p vector was generated similarly using a separate *Sac*II/*Sac*II product ligated into the *Sac*II site of YCplac111-MS2–Pab1p.

RBDs were determined according to the literature (**Table S1**) or using InterProScan software (Jones et al., 2014). Domains determined using InterProScan were extended by 30 residues on both ends. Primers were designed using Primer3Plus (Untergasser et al., 2007) with one additional nucleotide on both ends of the RBD (to maintain frame). Gateway B1 and B2 tails were included on the forward and reverse primers, respectively. Gateway reactions were performed as previously described (Walhout et al., 2000b).

### Yeast Manipulations and Assay Conditions

All assays were performed using the Y1H-aS2 yeast strain (Reece-Hoyes et al., 2011). Plasmid expressed baits were generated by yeast transformations as previously described (Walhout and Vidal, 2001) and plated on synthetic complete (Sc) -Trp agar media. Integrated baits were generated by transformation of yeast with *Nhe*I (NEB)-digested plasmids plated on Sc -His agar media. PRIMA assay strains were generated by yeast transformations of RNA-element harboring strains with individual prey plasmids plated on Sc -Leu, -Trp (plasmid baits) or Sc -Leu, -His (integrated baits). Individual colonies were picked and frozen at −80°C in 20% glycerol prior to performing the assay. All yeast strains are listed in **Table S2**.

Assays were performed as follows: Thawed yeast strains were inoculated in 200μl appropriate Sc liquid media in 96 deep well plates and grown overnight at 30°C with 200 rotations per minute (RPM) agitation. 10μl of overnight culture was diluted into 1mL of fresh media and grown to log phase (~6.5 h). Cultures were centrifuged at 2,000 RPM for 3 min. and resuspended in 400μl of 1X Phosphate Buffered Saline (PBS). Individual cells were then measured using a BD Accuri C6 flow cytometer using the 510/15 FL1 emission filter according to manufacturer's protocols.

### Data Processing and Quantitative Scoring

The standard flow cytometry data files (FCS3.0) were exported from BD Accuri C6 software and analyzed using custom R project software (http://www.R-project.org/) and the FlowCore and FlowViz packages. Briefly, forward scatter (FSC), side scatter (SSC) and fluorescence (FL1) measurements were imported for each sample. A lower FSC cutoff of 240,000 was applied as it corresponded to cellular debris (data not shown). A uniform cell population (~50% of the population) was selected using the FSC and SSC vectors and the norm2Filter function with scale factor=1. Briefly, the norm2filter function fits a bivariate normal distribution to the dataset and selects data points according to their standard deviation from the fit.

The resulting cells were plotted as fluorescence (FL1) *vs.* cell count and the two clear peaks were observed for nearly all cell populations. The low fluorescence peak overlapped with GFP-minus (LacZ) control yeast, indicating that zero GFP expression was present. The high fluorescence peak overlapped with GFP+ control yeast with poly(A) tails. We selected all ‘non-zero’ GFP cells by using a lower FL1 cutoff of 2048, which corresponded to the upper bound of GFP-control yeast. A FL1 cutoff of 1024 was used for the HBE:MS2BS RNA baits due to their low background. The population density was smoothed using a kernel density estimate. The peak of the density was determined for each sample. Eight replicates were tested for the initial experiments with the MS2BS, HBE, and RBP binding site baits (Figures 1 and 2). 16 replicates (two sets of eight) were collected for each 3′UTR bait and the two highest and two lowest values were removed. The average was calculated for the remaining 12 replicates from each bait-prey pair. The average fluorescence for each test prey was compared to the average SLBP-MS2-Pab1p negative control. Test preys with >1.20 fold increase in fluorescence were considered positive provided they were statistically significant (p<0.01, student's t-test).

### RNAi and Imaging of *C. elegans* Strains

Knockdowns were performed using the RNAi feeding method as described (Kamath et al., 2003). The RBD entry clones were cloned into the RNAi feeding vector construct L4440 using Gateway reactions and transformed into HT115(DE3) cells. The transformed colonies were grown to OD_600_ = 0.4 and induced with isopropyl 1-thio-β-D-galactopyranoside (IPTG) at a final concentration of 0.4mM for 4 hours. After induction the 50ml cultures were concentrated 10-fold and 50μl of the culture was added onto NGM plates containing 1mM IPTG and 100μg/ml Ampicillin. After bleaching adult animals in 0.5N NaOH and 2% clorox, eggs were washed once with distilled water, plated onto these plates and incubated at 25°C for 2 days before imaging. HT115 strain bacteria transformed with the empty vector L4440 was used as the control RNAi.

Adult animals were placed in 0.4mM levamisole on to 2% agarose pads before imaging. Embryo dissections were done in M9 solution and dissected eggs were mounted on 2% agarose pads. DIC and GFP fluorescence images were taken on Zeiss Axioscope 2 plus microscope (Carl Zeiss) using an oil-immersion 40X objective. Confocal images were taken under 40X magnification using Leica DMIRE2 microscope (Leica) using 488 nm excitation at 100% intensity. A single section was imaged for each worm and each line was scanned an average of 16 times to help eliminate background fluorescence.

## SUPPLEMENTAL INFORMATION

Supplementary Information includes four figures and two tables.

## AUTHOR CONTRIBUTIONS

A.M.T. and A.J.M.W. conceived the project, with critical advice from S.P.R and A.D.H. A.M.T. performed all cloning and yeast experiments with technical help from S.S. E.K. performed *C. elegans* experiments. A.M.T., A.D.H. and A.J.M.W. wrote the manuscript.

## ACKNOWLEDGEMENTS

The authors would like to thank Allan Jacobson, Job Dekker and members from the Walhout laboratory for advice and critical reading of the manuscript. Additionally, the authors would like to thank Allan Jacobson, Marvin Wickens and Paul Kaufman for reagents, and Phil Zamore and Nick Rhind for access to equipment. This work was supported by grants from the National Institutes of Health (HG006234 to A.J.M.W. and GM117237 to S.P.R.).

## REFFERENCES

Alberti, S., Gitler, A.D., and Lindquist, S. (2007). A suite of Gateway cloning vectors for high-throughput genetic analysis in Saccharomyces cerevisiae. Yeast 24, 913–919.

Amrani, N., Ganesan, R., Kervestin, S., Mangus, D.A., Ghosh, S., and Jacobson, A. (2004). A faux 3'-UTR promotes aberrant termination and triggers nonsense-mediated mRNA decay. Nature 432, 112–118.

Baltz, A.G., Munschauer, M., Schwanhausser, B., Vasile, A., Murakawa, Y., Schueler, M., Youngs, N., Penfold-Brown, D., Drew, K., Milek, M., et al. (2012). The mRNA-bound proteome and its global occupancy profile on protein-coding transcripts. Mol Cell 46, 674–690.

Bernstein, D., Hook, B., Hajarnavis, A., Opperman, L., and Wickens, M. (2005). Binding specificity and mRNA targets of a *C. elegans* PUF protein, FBF-1. RNA 11, 447–458.

Blazie, S.M., Babb, C., Wilky, H., Rawls, A., Park, J.G., and Mangone, M. (2015). Comparative RNA-Seq analysis reveals pervasive tissue-specific alternative polyadenylation in *Caenorhabditis elegans* intestine and muscles. BMC Biol 13, 4.

Butter, F., Scheibe, M., Morl, M., and Mann, M. (2009). Unbiased RNA-protein interaction screen by quantitative proteomics. Proc Natl Acad Sci U S A 106, 10626–10631.

Castello, A., Fischer, B., Eichelbaum, K., Horos, R., Beckmann, B.M., Strein, C., Davey, N.E., Humphreys, D.T., Preiss, T., Steinmetz, L.M., et al. (2012). Insights into RNA biology from an atlas of mammalian mRNA-binding proteins. Cell 149, 1393–1406.

Chalfie, M., Tu, Y., Euskirchen, G., Ward, W.W., and Prasher, D.C. (1994). Green Fluorescent Protein as a marker for gene expression. Science 263, 802–805.

Deplancke, B., Mukhopadhyay, A., Ao, W., Elewa, A.M., Grove, C.A., Martinez, N.J., Sequerra, R., Doucette-Stam, L., Reece-Hoyes, J.S., Hope, I.A., et al. (2006). A gene-centered *C. elegans* protein-DNA interaction network. Cell 125, 1193–1205.

Derti, A., Garrett-Engele, P., Macisaac, K.D., Stevens, R.C., Sriram, S., Chen, R., Rohl, C.A., Johnson, J.M., and Babak, T. (2012). A quantitative atlas of polyadenylation in five mammals. Genome Res 22, 1173–1183.

Dower, K., Kuperwasser, N., Merrikh, H., and Rosbash, M. (2004). A synthetic A tail rescues yeast nuclear accumulation of a ribozyme-terminated transcript. RNA 10, 1888–1899.

Farley, B.M., Pagano, J.M., and Ryder, S.P. (2008). RNA target specificity of the embryonic cell fate determinant POS-1. Rna 14, 2685–2697.

Farley, B.M., and Ryder, S.P. (2012). POS-1 and GLD-1 repress *glp-1* translation through a conserved binding-site cluster. Mol Biol Cell 23, 4473–4483.

Fuxman Bass, J.I., Sahni, N., Shrestha, S., Garcia-Gonzalez, A., Mori, A., Bhat, N., Yi, S., Hill, D.E., Vidal, M., and Walhout, A.J. (2015). Human gene-centered transcription factor networks for enhancers and disease variants. Cell 161, 661–673.

Gerstberger, S., Hafner, M., and Tuschl, T. (2014). A census of human RNA-binding proteins. Nat Rev Genet 15, 829–845.

Glisovic, T., Bachorik, J.L., Yong, J., and Dreyfuss, G. (2008). RNA-binding proteins and post-transcriptional gene regulation. FEBS Lett 582, 1977–1986.

Grove, C.A., deMasi, F., Barrasa, M.I., Newburger, D., Alkema, M.J., Bulyk, M.L., and Walhout, A.J. (2009). A multiparameter network reveals extensive divergence between *C. elegans* bHLH transcription factors. Cell 138, 314–327.

Hafner, M., Landthaler, M., Burger, L., Khorshid, M., Hausser, J., Berninger, P., Rothballer, A., Ascano, M., Jr., Jungkamp, A.C., Munschauer, M., et al. (2010). Transcriptome-wide identification of RNA-binding protein and microRNA target sites by PAR-CLIP. Cell 141, 129–141.

Hook, B., Bernstein, D., Zhang, B., and Wickens, M. (2005). RNA-protein interactions in the yeast three-hybrid system: affinity, sensitivity, and enhanced library screening. RNA 11, 227–233.

Hubstenberger, A., Cameron, C., Shtofman, R., Gutman, S., and Evans, T.C. (2012). A network of PUF proteins and Ras signaling promote mRNA repression and oogenesis in *C. elegans*. Dev Biol 366, 218–231.

Hunt-Newbury, R., Viveiros, R., Johnsen, R., Mah, A., Anastas, D., Fang, L., Halfnight, E., Lee, D., Lin, J., Lorch, A., et al. (2007). High-throughput in vivo analysis of gene expression in *Caenorhabditis elegans*. PLoS Biol 5, e237.

Jadhav, S., Rana, M., and Subramaniam, K. (2008). Multiple maternal proteins coordinate to restrict the translation of *C. elegans* nanos-2 to primordial germ cells. Development 135, 1803–1812.

Jan, C.H., Friedman, R.C., Ruby, J.G., and Bartel, D.P. (2011). Formation, regulation and evolution of Caenorhabditis elegans 3'UTRs. Nature 469, 97–101.

Johansson, H.E., Dertinger, D., LeCuyer, K.A., Behlen, L.S., Greef, C.H., and Uhlenbeck, O.C. (1998). A thermodynamic analysis of the sequence-specific binding of RNA by bacteriophage MS2 coat protein. Proc Natl Acad Sci U S A 95, 9244–9249.

Jones, P., Binns, D., Chang, H.Y., Fraser, M., Li, W., McAnulla, C., McWilliam, H., Maslen, J., Mitchell, A., Nuka, G., et al. (2014). InterProScan 5: genome-scale protein function classification. Bioinformatics 30, 1236–1240.

Kamath, R.S., Fraser, A.G., Dong, Y., Poulin, G., Durbin, R., Gotta, M., Kanapin, A., Le Bot, N., Moreno, S., Sohrmann, M., et al. (2003). Systematic functional analysis of the *Caenorhabditis elegans* genome using RNAi. Nature 421, 231–237.

Keene, J.D., Komisarow, J.M., and Friedersdorf, M.B. (2006). RIP-Chip: the isolation and identification of mRNAs, microRNAs and protein components of ribonucleoprotein complexes from cell extracts. Nat Protoc 1, 302–307.

Kemmeren, P., Sameith, K., van de Pasch, L.A., Benschop, J.J., Lenstra, T.L., Margaritis, T., O'Duibhir, E., Apweiler, E., van Wageningen, S., Ko, C.W., et al. (2014). Large-scale genetic perturbations reveal regulatory networks and an abundance of gene-specific repressors. Cell 157, 740–752.

Koh, Y.Y., Opperman, L., Stumpf, C., Mandan, A., Keles, S., and Wickens, M. (2009). A single *C. elegans* PUF protein binds RNA in multiple modes. RNA 15, 1090–1099.

Lee, I., Lehner, B., Crombie, C., Wong, W., Fraser, A.G., and Marcotte, E.M. (2008). A single gene network accurately predicts phenotypic effects of gene perturbation in *Caenorhabditis elegans*. Nat Genet 40, 181–188.

Li, S., Armstrong, C.M., Bertin, N., Ge, H., Milstein, S., Boxem, M., Vidalain, P.-O., Han, J.-D.J., Chesneau, A., Hao, T., et al. (2004). A map of the interactome network of the metazoan *C. elegans*. Science 303, 540–543.

Licatalosi, D.D., Mele, A., Fak, J.J., Ule, J., Kayikci, M., Chi, S.W., Clark, T.A., Schweitzer, A.C., Blume, J.E., Wang, X., et al. (2008). HITS-CLIP yields genome-wide insights into brain alternative RNA processing. Nature 456, 464–469.

Lublin, A.L., and Evans, T.C. (2007). The RNA binding proteins PUF-5, PUF-6, and PUF-7 reveal multiple systems for maternal mRNA regulation during *C. elegans* oogenesis. Dev Biol 303, 635–649.

MacNeil, L.T., Pons, C., Arda, H.E., Giese, G.E., Myers, C.L., and Walhout, A.J.M. (2015). Transcription factor activity mapping of a tissue-specific gene regulatory network. Cell Syst 1, 152–162.

Mangone, M., Manoharan, A.P., Thierry-Mieg, D., Thierry-Mieg, J., Han, T., Mackowiak, S., Mis, E., Zegar, C., Gutwein, M.R., Khivansara, V., et al. (2010). The Landscape of *C. elegans* 3'UTRs. Science 329, 432–435.

Mangus, D.A., Evans, M.C., and Jacobson, A. (2003). Poly(A)-binding proteins: multifunctional scaffolds for the post-transcriptional control of gene expression. Genome Biol 4, 223.

Martinez, N.J., Ow, M.C., Reece-Hoyes, J., Ambros, V., and Walhout, A.J. (2008). Genome-scale spatiotemporal analysis of *Caenorhabditis elegans* microRNA promoter activity. Genome Res 18, 2005–2015.

Matia-Gonzalez, A.M., Laing, E.E., and Gerber, A.P. (2015). Conserved mRNA-binding proteomes in eukaryotic organisms. Nat Struct Mol Biol 22, 1027–1033.

Merritt, C., Rasoloson, D., Ko, D., and Seydoux, G. (2008). 3'UTRs are the primary regulators of gene expression in the *C. elegans* germline. Curr Biol 18, 1476–1482.

Michel, F., Schumperli, D., and Muller, B. (2000). Specificities of *Caenorhabditis elegans* and human hairpin binding proteins for the first nucleotide in the histone mRNA hairpin loop. RNA 6, 1539–1550.

Moore, M.J. (2005). From birth to death: the complex lives of eukaryotic mRNAs. Science 309, 1514–1518.

Ogura, K., Kishimoto, N., Mitani, S., Gengyo-Ando, K., and Kohara, Y. (2003). Translational control of maternal *glp-1* mRNA by POS-1 and its interacting protein SPN-4 in *Caenorhabditis elegans*. Development 130, 2495–2503.

Okkema, P.G., Harrison, S.W., Plunger, V., Aryana, A., and Fire, A. (1993). Sequence requirements for myosin gene expression and regulation in *Caenorhabditis elegans*. Genetics 135, 385–404.

Opperman, L., Hook, B., DeFino, M., Bernstein, D.S., and Wickens, M. (2005). A single spacer nucleotide determines the specificities of two mRNA regulatory proteins. Nat Struct Mol Biol 12, 945–951.

Orr-Weaver, T.L., Szostak, J.W., and Rothstein, R.J. (1983). Genetic applications of yeast transformation with linear and gapped plasmids. Methods Enzymol 101, 228–245.

Pagano, J.M., Clingman, C.C., and Ryder, S.P. (2011). Quantitative approaches to monitor protein-nucleic acid interactions using fluorescent probes. RNA 17, 14–20.

Pagano, J.M., Farley, B.M., Essien, K.I., and Ryder, S.P. (2009). RNA recognition by the embryonic cell fate determinant and germline totipotency factor MEX-3. Proc Natl Acad Sci U S A 106, 20252–20257.

Pagano, J.M., Farley, B.M., McCoig, L.M., and Ryder, S.P. (2007). Molecular basis of RNA recognition by the embryonic polarity determinant MEX-5. J Biol Chem 282, 8883–8894.

Ray, D., Kazan, H., Chan, E.T., Pena Castillo, L., Chaudhry, S., Talukder, S., Blencowe, B.J., Morris, Q., and Hughes, T.R. (2009). Rapid and systematic analysis of the RNA recognition specificities of RNA-binding proteins. Nat Biotechnol 27, 667–670.

Reece-Hoyes, J.S., Diallo, A., Kent, A., Shrestha, S., Kadreppa, S., Pesyna, C., Lajoie, B., Dekker, J., Myers, C.L., and Walhout, A.J.M. (2011). Enhanced yeast one-hybrid (eY1H) assays for high-throughput gene-centered regulatory network mapping. Nature Methods 8, 1059–1064.

Reece-Hoyes, J.S., Pons, C., Diallo, A., Mori, A., Shrestha, S., Kadreppa, S., Nelson, J., DiPrima, S., Dricot, A., Lajoie, B.R., et al. (2013). Extensive rewiring and complex evolutionary dynamics in a C. elegans multiparameter transcription factor network. Mol Cell 51, 116–127.

Ritter, A.D., Shen, Y., Bass, J.F., Jeyaraj, S., Deplancke, B., Mukhopadhyay, A., Xu, J., Driscoll, M., Tissenbaum, H.A., and Walhout, A.J. (2013). Complex expression dynamics and robustness in *C. elegans* insulin networks. Genome research 23, 954–965.

SenGupta, D.J., Zhang, B., Kraemer, B., Pochart, P., Fields, S., and Wickens, M. (1996). A three-hybrid system to detect RNA-protein interactions in vivo. Proc Natl Acad Sci U S A 93, 8496–8501.

Stellberger, T., Hauser, R., Baiker, A., Pothineni, V.R., Haas, J., and Uetz, P. (2010). Improving the yeast two-hybrid system with permutated fusions proteins: the Varicella Zoster Virus interactome. Proteome science 8, 8.

Stumpf, C.R., Kimble, J., and Wickens, M. (2008). A *Caenorhabditis elegans* PUF protein family with distinct RNA binding specificity. RNA 14, 1550–1557.

Szostak, E., and Gebauer, F. (2013). Translational control by 3'-UTR-binding proteins. Briefings in functional genomics 12, 58–65.

Tamburino, A.M., Ryder, S.P., and Walhout, A.J. (2013). A compendium of *Caenorhabditis elegans* RNA binding proteins predicts extensive regulation at multiple levels. G3 (Bethesda) 3, 297–304.

Tenenbaum, S.A., Carson, C.C., Lager, P.J., and Keene, J.D. (2000). Identifying mRNA subsets in messenger ribonucleoprotein complexes by using cDNA arrays. Proc Natl Acad Sci U S A 97, 14085–14090.

Thompson, B.E., Bernstein, D.S., Bachorik, J.L., Petcherski, A.G., Wickens, M., and Kimble, J. (2005). Dose-dependent control of proliferation and sperm specification by FOG-1/CPEB. Development 132, 3471–3481.

Ule, J., Jensen, K., Mele, A., and Darnell, R.B. (2005). CLIP: a method for identifying protein-RNA interaction sites in living cells. Methods 37, 376–386.

Ulitsky, I., Shkumatava, A., Jan, C.H., Subtelny, A.O., Koppstein, D., Bell, G.W., Sive, H., and Bartel, D.P. (2012). Extensive alternative polyadenylation during zebrafish development. Genome Res 22, 2054–2066.

Untergasser, A., Nijveen, H., Rao, X., Bisseling, T., Geurts, R., and Leunissen, J.A. (2007). Primer3Plus, an enhanced web interface to Primer3. Nucleic Acids Res 35, W71–74.

Walhout, A.J.M. (2011). What does biologically meaningful mean? A perspective on gene regulatory network validation. Genome Biol 12, 109.

Walhout, A.J.M., Sordella, R., Lu, X., Hartley, J.L., Temple, G.F., Brasch, M.A., Thierry-Mieg, N., and Vidal, M. (2000a). Protein interaction mapping in *C. elegans* using proteins involved in vulval development. Science 287, 116–122.

Walhout, A.J.M., Temple, G.F., Brasch, M.A., Hartley, J.L., Lorson, M.A., van den Heuvel, S., and Vidal, M. (2000b). GATEWAY recombinational cloning: application to the cloning of large numbers of open reading frames or ORFeomes. Methods in enzymology: "Chimeric genes and proteins" 328, 575–592.

Walhout, A.J.M., and Vidal, M. (2001). High-throughput yeast two-hybrid assays for large-scale protein interaction mapping. Methods 24, 297–306.

Wang, X., Zhao, Y., Wong, K., Ehlers, P., Kohara, Y., Jones, S.J., Marra, M.A., Holt, R.A., Moerman, D.G., and Hansen, D. (2009). Identification of genes expressed in the hermaphrodite germ line of C. elegans using SAGE. BMC Genomics 10, 213.

Watson, E., MacNeil, L.T., Arda, H.E., Zhu, L.J., and Walhout, A.J.M. (2013). Integration of metabolic and gene regulatory networks modulates the *C. elegans* dietary response. Cell 153, 253–266.

Yu, H., Tardivo, L., Tam, S., Weiner, E., Gebreab, F., Fan, C., Svrzikapa, N., Hirozane-Kishikawa, T., Rietman, E., Yang, X., et al. (2011). Next-generation sequencing to generate interactome datasets. Nat Methods 8, 478–480.

Zhang, B., Kraemer, B., SenGupta, D., Fields, S., and Wickens, M. (1999). Yeast three-hybrid system to detect and analyze interactions between RNA and protein. Methods Enzymol 306, 93–113.

